# Using Chaos for Facile High-throughput Fabrication of Ordered Multilayer Micro- and Nanostructures

**DOI:** 10.1101/833772

**Authors:** Carolina Chávez-Madero, María Díaz de León-Derby, Mohamadmahdi Samandari, Carlos Fernando Ceballos-González, Edna Johana Bolívar-Monsalve, Christian Carlos Mendoza-Buenrostro, Sunshine Holmberg, Norma Alicia Garza-Flores, Mohammad Ali Almajhadi, Ivonne González-Gamboa, Juan Felipe Yee-de León, Sergio Omar Martínez-Chapa, Ciro A. Rodríguez, Hemantha Kumar Wickramasinghe, Marc Madou, Ali Khademhosseini, Yu Shrike Zhang, Mario Moisés Álvarez, Grissel Trujillo-de Santiago

## Abstract

This paper introduces the concept of continuous chaotic printing, i.e., the use of chaotic flows for deterministic and continuous fabrication of fibers with internal multilayered micro-or nanostructures. Two free-flowing materials are coextruded through a printhead containing a miniaturized Kenics static mixer (KSM) composed of multiple helicoidal elements. This produces a fiber with a well-defined internal multilayer microarchitecture at high speeds (>1.0 m min-1). The number of mixing elements and the printhead diameter determine the number and thickness of the internal lamellae, which are generated according to successive bifurcations that yield a vast amount of inter-material surface area (~102 cm2 cm3) and high resolution features (~10 μm). In an exciting further development, we demonstrate a scale-down of the microstructure by 3 orders of magnitude, to the nanoscale level (~10 nm), by feeding the output of a continuous chaotic 3D printhead into an electrospinner. Comparison of experimental and computational results demonstrates the robust and predictable output and performance of continuous chaotic 3D printing. The simplicity and high resolution of continuous chaotic printing strongly supports its potential use in novel applications, including—but not limited to—bioprinting of multi-scale tissue-like structures, modeling of bacterial communities, and fabrication of smart multi-material and multilayered constructs.

## Introduction

In nature and engineering, multi-material and multi-layered architectures achieve functionality or performance that is not achievable with monolithic materials. Moreover, the functionality and performance of multilayered composites is frequently determined by the degree of intimacy, or the density of their layers. Multilayered materials with a high amount of inter-material area can yield higher capacitances in supercapacitors,^[1]^ elevated mechanical strength^[2,3]^ and fatigue resistance,^[4]^ better sensing capabilities,^[5]^ or improved energy-harvesting potential.^[6]^ A multi-lamellar architecture that features a tight control of the surface area is also desirable in applications related to the controlled release of pharmaceuticals.^[7]^ Although appealing and enabling, the cost-effective fabrication of multi-material lamellar microarchitecture has proven to be challenging. Some of the current micro-and nano-manufacturing technologies such as stereolithography, three-dimensional (3D) printing, and nano-imprinting are capable of fabricating relatively complex lamellar architectures, but often require a costly investment in terms of equipment, skilled labor, and time. These technologies typically require using optimized inks, a specific and narrow range of conditions and normally involve long fabrication times and accurate control systems.^[8,9]^ Multi-material 3D printing has enabled the use of multiple inks in the same printing operation, but faces severe limitations in resolution and speed.^[10,11]^ A combination of multiple channels, each one dispensing one material, has been demonstrated to fabricate multi-material constructs with resolutions in the range of 50-100 micrometers.^[12–17]^ We recently demonstrated multi-material 3D printing of perusable multi-layered cannulas by co-extruding multiple streams of inks through a set of concentric capillary tubes contained in a single nozzle.^[18]^ Similarly, Kang and coworkers presented an extrusion printing technique that produced multi-material tissue-like microstructures by coextrusion of different materials through a head with a pre-set internal architecture.^[16]^ These state-of-the-art approaches to 3D printing produce structures at resolutions dictated (in the best scenario) by the smallest relevant length scale of the nozzle^[10]^ and exhibit only moderate speeds. To date, no study has demonstrated the robust, fast, and cost-effective fabrication of reproducible micro-and nanostructures in a multi-material construct through a single-nozzle printhead.

## Results and Discussion

### Continuous chaotic printing: A simple and effective microfabrication strategy

Here we introduce the concept of continuous chaotic printing: the use of a simple laminar chaotic flow induced by a static mixer for the continuous creation of fine and complex structures at the micrometer and submicrometer levels within polymer fibers. Chaotic flows are used to mix in the laminar regime, where the conditions of low speed and high viscosity preclude the use of turbulence to achieve homogeneity.^[19,20]^ In the context of 3D printing, they have been suggested as a tool to provide better homogenization of different materials.^[21,22]^ However, a much less exploited characteristic of chaotic flows is their potential to create defined multi-material and multi-lamellar structures.^[23–26]^

A recent contribution from our group demonstrated, for the first time, the use of simple chaotic flows (i.e., Journal Bearing flow) to imprint fine microstructures within constructs in a controlled and predictable manner at an exponentially fast rate in a batch-wise fashion.^[27]^ In the present study, we explore the utility of a chaotic printer, equipped with a Kenics static mixer (KSM)^[28]^ as a key component of the printhead, for the printing of fibers with massive amounts of lamellar microstructures in a continuous fashion (**Figure 1**).

**Figure 1.**
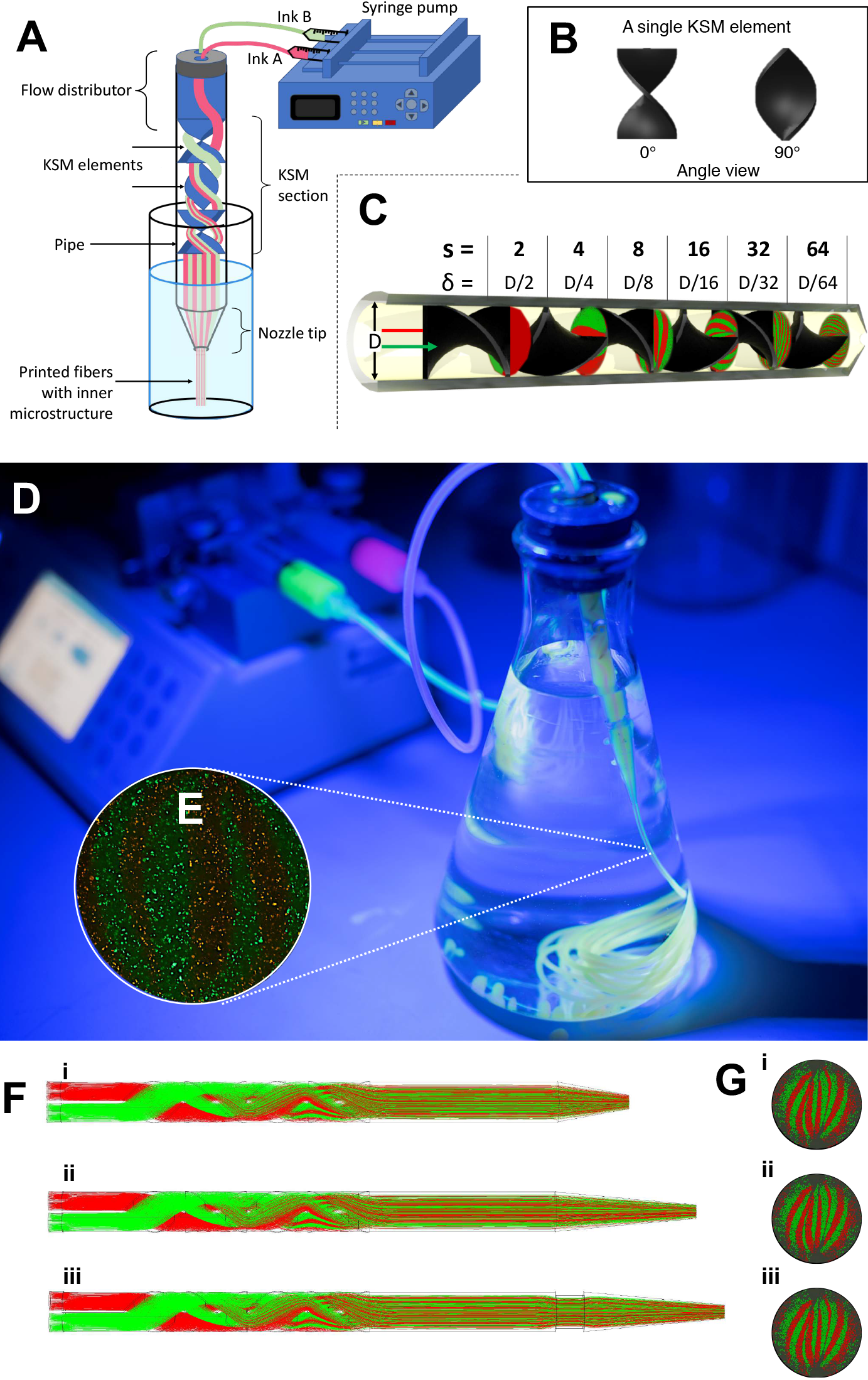
Experimental setup. Continuous chaotic printing is based on the ability of a static mixer to create structure within a fluid. The Kenics static mixer (KSM) induces a chaotic flow by a repeated process of reorientation and splitting of fluid as it passes through the mixing elements. (A) Schematic representation of a KSM with two inlets on the lid. The inks are fed at a constant rate through the inlets using syringe pumps. The inks flow across the static mixer to produce a lamellar structure at the outlet. The inks are crosslinked at the exit of the KSM to stabilize the structure. Our KSM design includes a cap with 2 inlet ports, a straight non-mixing section that keeps the ink injections independent, a mixing section containing one or more mixing elements, and a nozzle tip. The lid can be adapted to inject several inks simultaneously. (B) Two rotated views at 0° and 90°, of a single KSM element. (C) 3D design of a KSM with 6 elements and schematic representation of the flow splitting action, the increase in the number of striations, and the reduction in length scales, in a KSM-printhead. The resolution, namely the number of lamellae and the distance between them (δ), can be tuned using different numbers of KSM elements. (D) Actual continuous chaotic printing in operation. The inset (E) shows the inner lamellar structure formed at the cross-section of the printed fiber (the use of 3 KSM elements originates 8 striations). (F) Longitudinal or (G) cross-sectional microstructure of fiber obtained using different tip nozzle geometries. Images show CFD results of particle tracking experiments where two different inks containing red or green particles are coextruded through a printhead containing 4 KSM elements. The lamellar structure is preserved when the outlet diameter is reduced, from 4 mm (inner diameter of the pipe section) to 2 mm (inner diameter of the tip), through tips differing in their reduction slope.

Our chaotic printer is composed of a flow distributor, a pipe, a static mixing section, and an outlet or nozzle tip (Figure 1A,B). The number of inks one can use is unrestricted, and different distributor geometries can be employed to accommodate the injection of multiple inks. However, in this communication we adhered to some of the simplest printing scenarios. To this end, we adopted a distributor configuration (Figure 1A,C) for dispensing two inks in a symmetrical fashion. The mixing section contains a KSM, a static mixer configuration widely adopted in the chemical industry, that consists of a serial arrangement of *n* number of helical elements contained in a tubular pipe, with each element is rotated 90° with respect to the previous one (Figure 1B,C). In the laminar regime, the KSM (and other static mixers^[29–31]^) produces chaos by repeatedly splitting and reorienting materials as they flow through each element. With this simple mechanism, lamellar interfaces are effectively produced between fluids (i.e., printheads containing 1, 2, 3, 4, 5, or 6 KSM elements will produce 2, 4, 8, 16, 32, or 64 defined striations; Figure 1C; Figure S1). Our results show that multimaterial lamellar structures with different degrees of inter-material surface can be printed using a single nozzle by simply co-extruding two different materials (i.e., inks) through a KSM.

In the experiments presented here, we used sodium alginate to formulate different inks consisting of pristine alginate or suspensions of particles (polymer microparticles, graphite microparticles, mammalian cells or bacteria). For instance, we conducted experiments in which one (i.e., red bacteria) or two type(s) of fluorescent microparticles (i.e., red and green bacteria or polymer beads) were injected into the ports of the mixer distributor (Figure 1D,E). The result is continuous composite fibers with complex microstructures that can be stabilized simply by crosslinking in a bath of calcium chloride solution, which preserved the internal microstructure of the fibers with high fidelity (Figure 1D,E; Figure S1). Fine and well-aligned microstructures with defined features can be robustly fabricated along the printed fibers at remarkably high extrusion speeds (1–5 m of fiber/min). As we will show later, a vast amount of contact area is developed within each linear meter of these fibers. This printing strategy is also robust across a wide range of operation settings. We conducted a series of printing experiments at different inlet flow rates to assess the stability of the printing process. As long as the flow regime is laminar and the fluid behaves in a Newtonian manner (Figure S2), the quality of the printing process is not affected by the flow rate used in a wide range of flow conditions. For example, using a cone-shaped nozzle-tip with an outlet diameter of 1 mm, stable fibers were obtained in a window of flow rates from 0.003 to 5.0 mL/min (Figure 1D). Having printheads with different geometries (different degrees of slope) did not disturb the lamellar structure generated by chaotic printing. CFD simulation results suggested that the angle of inclination of the conical tip of the printhead (nozzle tip) did not appear to significantly affect the microstructure within the fiber in the range of tested flow rates and reduction slopes. Figure 1F,G shows a computational analysis of the effect of the shape of the printhead tip (angle) on the conservation of the microstructure of printed fibers produced from a mixture of alginate inks containing red and green particles.

### Coupling of continuous chaotic printing with other fabrication techniques

The combination of continuous chaotic printing with other fabrication technologies (e.g., molding, electrospinning, or robotic assembly) will lead to the fabrication of complex multi-scale architectures with high degrees of predictable external shapes and internal microstructure. Indeed, during printing, these fibers can be rearranged either into macrostructures or the individual fibers can be further reduced in diameter while preserving their lamellar architecture (Figure 1F,G: Figure 2a-c). We illustrate the former by printing a long fiber of alginate containing multiple lamellae and then rearranging it into a block of several layers of fiber segments (Figure 2A-C). The integration of this multi-material printhead into a 3D printer may thus enable rapid fabrication of multi-material (and/or multi-cellular) constructs that exhibit a great amount of material interface with a complex and tunable architecture as we will show later in this contribution. Also, to demonstrate the latter (reducing the fiber diameter), chaotic printing may be coupled with other techniques for the production of nanofibers that contain finely controlled structures at the submicron scale (Figure 2D-G).

**Figure 2.**
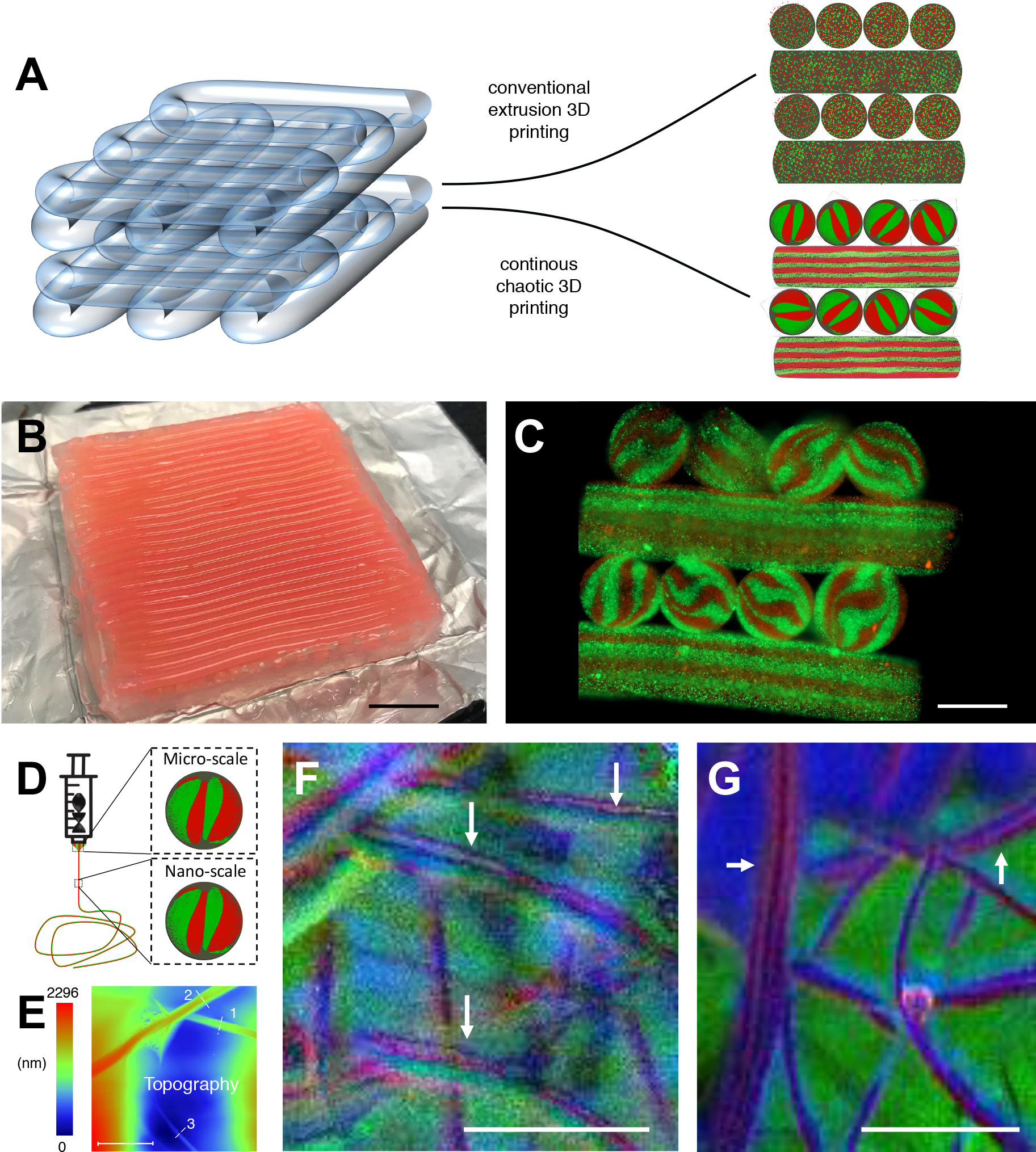
Development of multi-scale architectures based on 3D continuous chaotic printing: (A-C) 3D printing of hydrogel constructs using a KSM-printhead integrated to a commercial cartesian 3D printer. (A) Schematic comparison of the lack (prepared using conventional extrusion techniques) and presence of internal lamellar microstructures (developed using continuous chaotic printing). (B) Printing of a long fiber arranged into a macro-scale hydrogel construct (3 cm × 3 cm × 4 mm). Scale bar: 5 mm. (C) Transverse cut of the macro-construct showing the internal microstructures. Scale bar: 1 mm. (D-G) Chaotic printing of fibers coupled with electrospinning. (D) Schematic representation of the coupling between continuous chaotic printing and an electrospinning platform; an ink composed of a pristine alginate ink (4% sodium alginate in water) and an ink composed of a polyethylene oxide blend (7% polyethylene oxide in water), were coextruded through a chaotic printhead and electrospun into a nanomesh. (E) AFM image showing the diameter of three individual nanofibers [(1) 0.82 μm, (2) 1.05 μm, and (3) 0.437 μm] within the electrospun mesh. Scale bar: 5 μm. (F,G) PIFM reveals the lamellar nature of the nanostructure within a nanofiber (white arrows) originated using (F) a 2-element KSM printhead, and (G) a 3-element KSM printhead. Scale bar: 1 μm.

As an example, we coupled a 2-element KSM printhead with an electrospinning device (Figure 2D) to produce a mesh of nanofibers containing well-defined nanostructures. Fibers of a mean diameter <300 nm were formed (Figure 2E). A close inspection using photo-induced force microscopy (PIFM)^[32]^ revealed multilayered nanostructures with average striation thicknesses in the range of 75–100 nm (Figure 2F,G; Figure S3). These results demonstrated that the microstructure created by 3D chaotic printing can be further scaled down by three orders of magnitude using electrospinning. The fabrication of fibers with fine lamellar microstructures will enable the design of materials with many relevant applications. Next, we discuss exemplary applications of continuous chaotic printing for the rapid creation of lamellar interfaces in the context of material sciences and biology.

### Multilayered and well-aligned microstructures

In Figure 3, we present the results of an experiment in which a suspension of 0.5% graphite microparticles in pristine alginate ink (1%) was co-extruded with pristine alginate ink (1%). For this experiment, the printhead outlet had a diameter of 2 mm (Figure 3A). Note that the features in the extruded structure were remarkably similar at different lengths of the fiber (Figure 3B), demonstrating the robustness of this printing strategy with small nozzle diameters.

**Figure 3.**
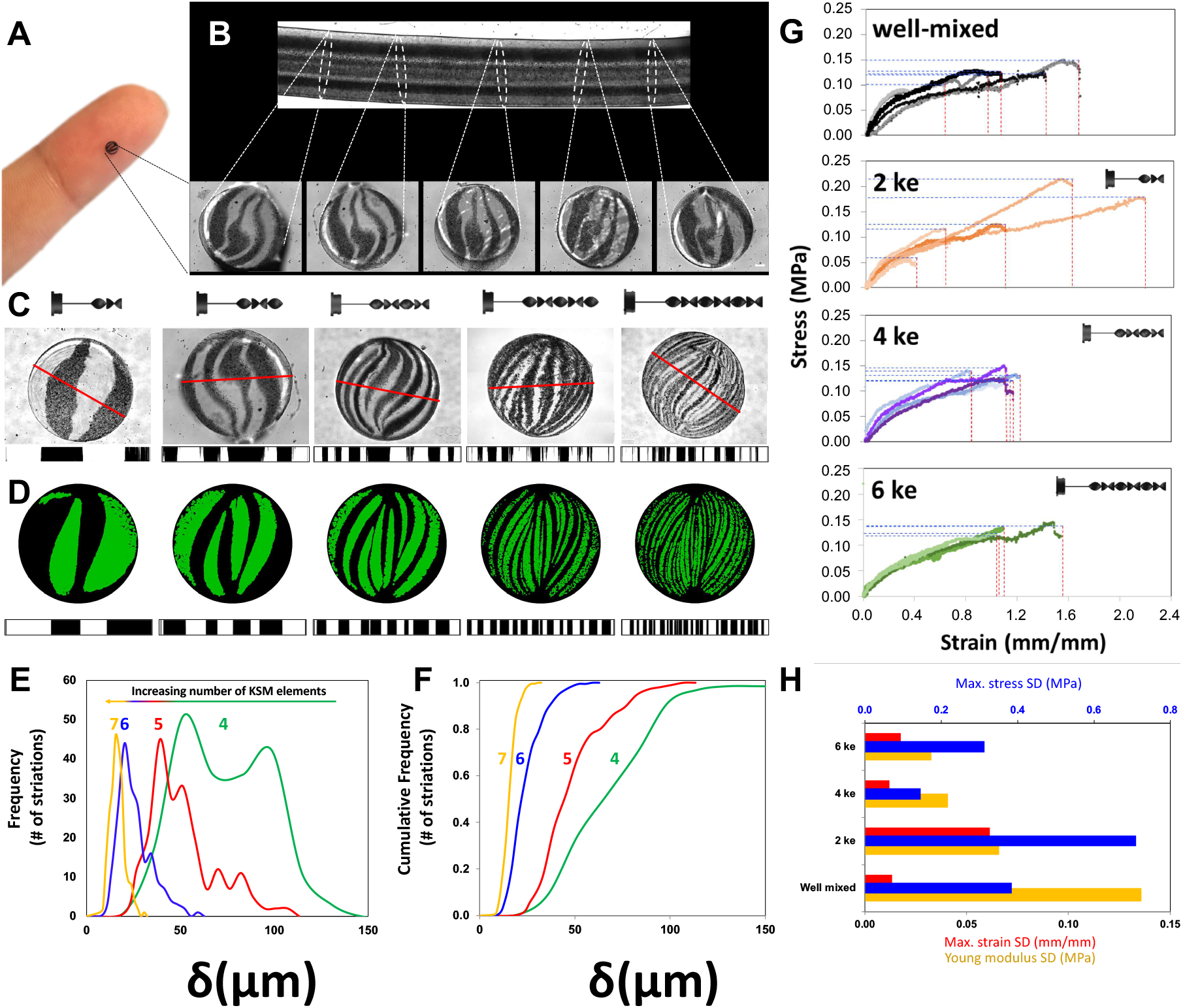
Evaluation of the striation profiles and mechanical properties of chaotically printed alginate/graphite fibers. (A, B) Cross-sectional cuts along an alginate/graphite fiber printed using 3 KSM elements. Scale bar: 200μm. (C) Lamellar microstructure of fibers produced with printheads containing 2, 3, 4, 5, or 6 KSM elements. The thickness of each lamella, along the red line, was determined by image analysis using Image J (shown below each cross-sectional cut). Scale bar (red): 2 mm. (D) The microstructure at each cross-section was reproduced by CFD simulations, and the thickness and position of each lamella was calculated. (E) Striation Thickness Distribution (STD) and (F) cumulative STD for constructs printed using 4, 5, 6, and 7 KSM elements. (G) Comparison of stress-strain curves of fibers fabricated by extrusion of pristine alginate and graphite without chaotic mixing (marked as hand-mixed) or with chaotic printing using 2, 4, or 6 KSM elements (marked as 2, 4, or 6). (H) Comparison of the standard deviation of tensile properties (i.e., maximum stress, maximum strain, and Young’s modulus for the same set of fibers; 5 fibers per treatment).

Notably, the resolution of this technique is controlled by both the diameter of the nozzle and the number of mixing elements. As the number of elements used to print increased, the number of lamellae observed in any given cross-sectional plane of the fiber also increased, while the thickness of each lamella decreased (Figure 3C).

Therefore, users of continuous chaotic printing will have more degrees of freedom to determine the multi-scale resolution of a construct, as this is no longer mainly restricted by the diameter of the nozzle (or the smallest length-scale of the nozzle at cross-section). For instance, for our two-stream system (Figure 1C), the number of lamellae increases exponentially according to the simple model *s*=2^*n*^, where *s* is the number of lamellae or striations within the construct and *n* is the number of KSM elements within the extrusion tube. Two streams of inks co-injected into the printhead will generate 4, 8, 16, 32, 64, and 128 distinctive streams of fluid when passing through a series of 2, 3, 4, 5, 6, and 7 KSM elements, respectively (Figure 3C). The average resolution of the structure will then be governed by the average striation of the construct (δ), given by δ =D/s, were D is the nozzle inner diameter. Since stretching is exponential in chaotic flows,^[23, 25,27]^ the reduction in the length scale is also exponential, as is the increase in resolution (i.e., more closely packed lines; see also Figure S1). In the experiment portrayed in Figure 3, the cross-sectional diameter of the fibers was 2mm. We observed defined average striations with resolutions of ~500, 250, 125, 62.5, and 31.75 μm by continuously printing using 2, 3, 4, 5, and 6 KSM elements, respectively. Even when 6 KSM elements were used, distinctive lamellae could be discriminated in the array of 64 aligned striations (Figure 2C). The resolution values obtained through 6 elements already exceeded those achievable by state-of-the-art commercial 3D extrusion printers (~100–75 μm)^[33,34]^ that use hydrogel-based inks (i.e., commercial bioprinters).^[33,35]^

Another remarkable characteristic of continuous chaotic printing is that the structure obtained is fully predictable, since chaotic flows are deterministic systems (as in any chaotic system).^[25,36]^ Simulation results, obtained by solving the Navier-Stoke equations of fluid motion using computational fluid dynamics (CFD),^[37,38]^ closely reproduced the cross-sectional lamellar microarchitecture within the fibers (Figure 1F,G; Figure 3D).

Moreover, we used optical microscopy and image analysis techniques to characterize the fine array of lamellae experimentally produced by continuous chaotic printing. We calculated the striation thickness distribution (STD) on the cross-sections of the graphite/alginate fibers. We did this by drawing several center lines of representative cross-sections and then calculating the distance between striations along those lines (Figure S4). The frequency distribution and the cumulative STD were then measured. Figure 3e and f, respectively, show the STD and the cumulative STD for constructs printed using 4, 5, 6, and 7 KSM elements. Remarkably, this family of distributions exhibits self-similarity, one of the distinctive features of chaotic processes.^[24,25,27]^ As discussed, for any of these particular cases, the average striation thickness could be calculated as the fiber diameter/number of striations. However, due to the highly skewed shape of the distribution toward smaller striation thicknesses (Figure 3F), the median striation thickness is lower than the average striation value. For example, for the case where 4 KSM elements were used, the average striation thickness can be calculated as 2 mm/16 = 125 μm. Indeed, 50% of the striations measured less than 125 μm (Figure 3E,F), but the corresponding STD showed that most of the striations had a median value of about 75 μm. This has profound implications for crucial processes such as mass and heat transfer. Remarkably, the diffusional distances in these constructs decreased rapidly as the number of elements used to print was increased.

Fiber and particle alignment is key in many applications in materials technology.^[39]^ We next show that the fabrication of well-aligned microstructures achievable through chaotic printing can influence relevant characteristics of composites, such as the robustness of their mechanical performance. We characterized the mechanical properties of fibers 2.5 cm in length produced by chaotic printing (and therefore having different internal structures) or random mixing and extrusion through an empty pipe (Figure 3; Figure S5). Specifically, we conducted tensile testing using a universal testing machine on fibers produced from a mixture of 0.5% graphite microparticles in alginate, generated either by random mixing (a control without lamellar structures) or by continuous chaotic printing using 2, 4, or 6 KSM elements. Figure 3G shows the stress-strain curves associated with the resulting fibers. We did not find significant differences in the Young’s modulus, ultimate stress, or maximum elongation at break in these sets of fibers (Figure S5). However, the fibers exhibited less variability when produced by continuous chaotic printing than those by extrusion of randomly mixed inks. Among the fibers produced by chaotic printing, the fibers were more homogeneous when co-extruded through printheads containing 4 or 6 elements than only 2 elements. Figure 2H shows an analysis of the standard deviation of relevant mechanical performance indicators associated with different microstructures. These results suggest that effective alignment of the microstructures within the fibers resulted in a more reproducible mechanical performance.

### Bioprinting applications

Bioprinting (i.e., the printing of cells, proteins, and biomaterials in a predefined fashion) is presently even more limited in resolution and speed than additive manufacturing techniques in general. We further illustrate a biological application of continuous chaotic printing by fabricating constructs with specific microarchitectures containing living cells.

Tightly controlling the degree of intimacy (i.e., the density of interfaces between bacterial populations) may enable the fabrication of 3D multi-material constructs with novel functionalities^[40]^ and is of paramount importance in modern microbiology.^[41]^ The spatial arrangement and distribution of bacteria, recently described as “microbiogeography”,^[41]^ is an important determinant of bacterial community dynamics, as different species of bacteria interact with other microorganisms through chemical signals.^[41–43]^ Bacterial chemotaxis (i.e., quorum sensing (QS)), a well-studied phenomenon, depends on the vicinity and the amount of surface area shared among bacterial communities.^[44]^ The efficacy of QS, one of the mechanisms that plays a key role in bacterial virulence, is known to be influenced by distance and spatial distribution,^[45,46]^ but relatively few studies have addressed the relationship between spatial distribution, distance, and cell density in bacterial systems.^[45–47]^ Continuous chaotic printing enables precise control of the spatial distribution of bacterial communities, aligned in a micro-lamellar microstructure, and allows meticulous design and regulation of the amount of interface between bands of bacteria.

In Figure 4A-E, we used two recombinant *Escherichia coli* strains, one producing red fluorescent protein (RFP) and the other producing green fluorescent protein (GFP) to fabricate cell-laden fibers. As anticipated, well-defined bacterial striations could be printed by our technique (Figure 4A). Remarkably, the bacteria could be cultured in these fibers for extended time periods. We followed the kinetic behavior of both bacterial populations (i.e., GFP-and RFP-bacteria) in the fibers initially seeded at low concentrations. During the first 24 h of culture (from t = 0 to 24 h), the intensity of the fluorescence produced by the bacterial colonies increased, while the bacteria continued to respect the original patterns in which they had been printed (Figure 4B).

**Figure 4.**
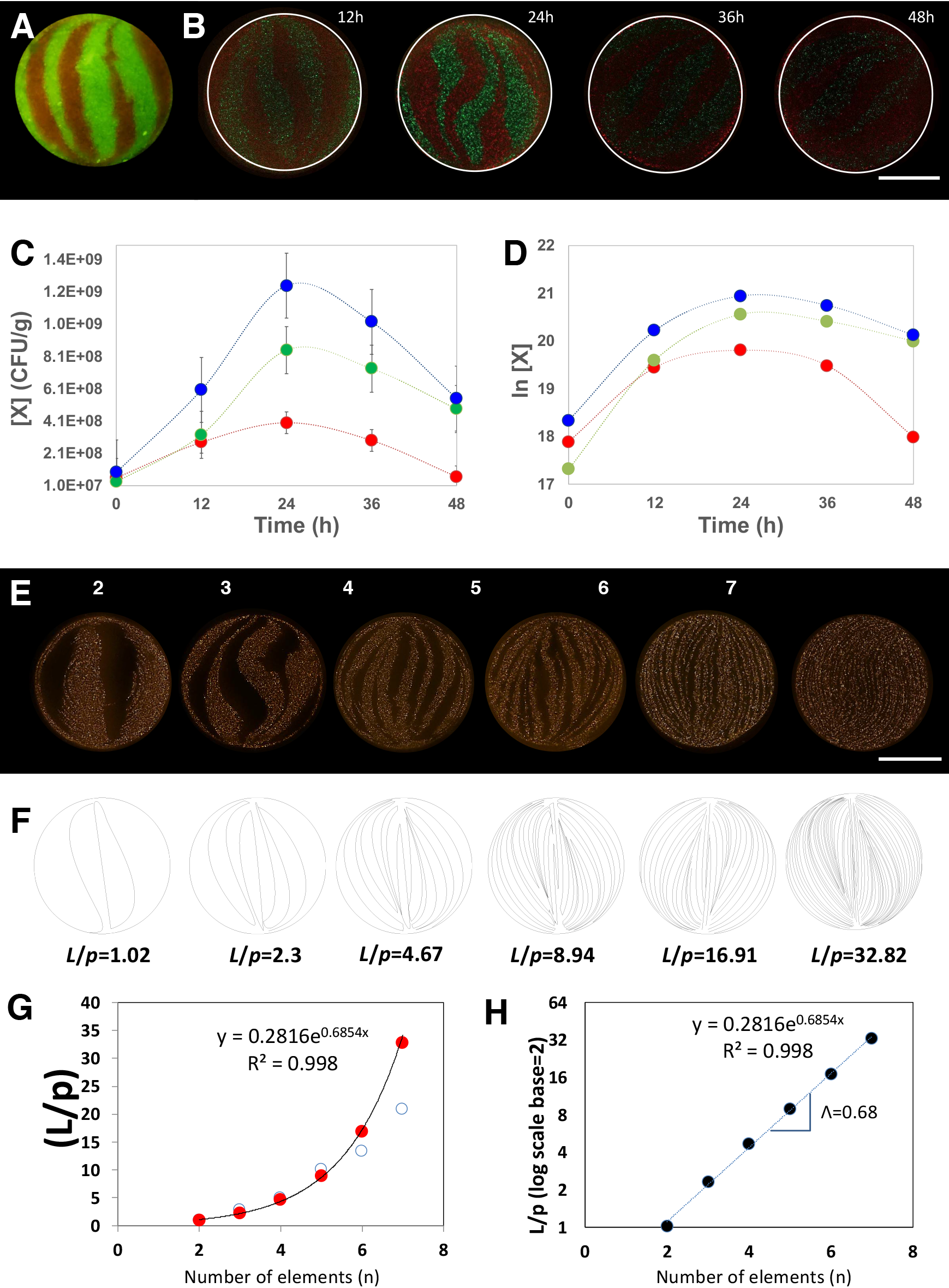
Chaotic bioprinting of bacteria. (A) Cross-section of a fiber where GFP-and RFP-bacteria shared an inter-material interface. The micrograph was obtained after chaotic printing at a high initial cell concentration and using 3 KSM elements. Scale bar: 250 μm. (B) The evolution of the concentration of living bacteria in the cross section of a fiber, initially printed with a low bacterial concentration, shown by micrographs taken at 0, 12, 24, and 48 h; Scale bar: 250 μm. (C) Growth curves showing the increasing concentration of viable cells over time, as determined by standard plate culture microbiological methods. Red and green symbols indicate the evolution of red and green fluorescent bacteria, respectively. Blue symbols show the total numbers of viable cells. (D) Plot of the natural log of bacterial populations over time. (E) Cross-sections of alginate fibers containing fine and aligned striations of RFP-*E. coli*. These fibers of 1 mm thickness were produced by chaotic bioprinting using printheads containing 2 to 7 KSM elements. Scale bar: 250μm. (F) Determination of the shared interface from computational simulations. (G) Estimation of the total amount of interface shared between regions with and without bacteria (*L*), normalized by the perimeter of the fiber (*p*); red dots (o) indicate approximations based on a simple geometric model, and empty blue circles (•) show determinations based in image analysis of experimentally obtained micrographs. (H) Natural log of the *L*/*p* ratio as a function of the number of elements used to print. The Lyapunov exponent (Λ) of the chaotic flow is calculated from the slope of the resulting straight line.

We corroborated the increase in the number of live bacteria by conventional colony-forming units (CFU) microbiological assays (Figure 4C,D). To do this, we sampled multiple sections of fibers. We consistently observed that the bacterial populations grew in both areas for the first 24 h, exhibited a short plateau, and then later decayed. These results demonstrated that chaotic printing can be used for the fabrication of dynamic living systems that are capable of evolving in time from very well-defined initial conditions. In addition, massive amounts of interface between green and red bacterial regions could be developed if more elements were used during printing.

For example, the boundary between the green and red bacterial regions can be effectively tuned from ~1 mm down to 15 μm by varying the number of KSM elements used to print (from 2 to 7, Figure 4E). Since the maximum length of these bacteria is ~2 μm, we were able to imprint lamellae of bacteria in the resolution range of tens of micrometers. The diameter of these fibers was 1 mm. Therefore, printing using 7 KSM elements yielded lamellae with average striation thicknesses of less than 10 μm (median lower than 7 μm). This means that each lamella might accommodate a few bacterial cells across its width.

In chaotic printing, the amount of inter-material area fabricated increases exponentially as a function of the number of elements used. We used image analysis techniques to quantify, at high magnification, the shared perimeters between red and black lamellae in Figure 4E (cross-sectional cuts). Indeed, the amount of black-red perimeter at cross-sections grew exponentially with the increasing number of elements (Figure 4E,G).

In addition, using computational strategies, we simulated the amount of surface area generated by the printing process in constructs printed using different KSM elements (Figure 4F), and confirmed the data obtained experimentally (Figure S6; Figure 4G). For instance, when 6 elements were used to print, approximately 15 cm of shared linear interface were developed between the two materials (inks) at each cross-sectional plane (D = 0.1 cm); the ratio between the total amount of developed interface and the fiber perimeter was 16.91. This created a remarkably high density of shared interface (6.76 cm/mm^2^). Since the fiber exhibits the very same microstructure along its entire length (Figure 3B, Figure S1B), the inter-material surface density generated inside the fiber could be determined as ~0.067 m^2^ cm^−3^. For example, using a printhead containing 6 KSM elements, 0.067 m^2^ of material interface could be accommodated per 1 cm^3^ of living fiber. Therefore, ~67 m^2^ L^−1^ of well-aligned inter-material surfaces could be fabricated within these living constructs.

For comparison, human kidneys have an approximate volume of 150 cm^3^, and the total area of the capillaries of all the glomeruli within them is 0.6 m^2^ (4 m^2^ L^−1^).^[48]^ Printing at the flow rate of 1 mL/min, which is a typical printing flow rate used in our system, could generate this amount of area per unit of volume every minute. This massive amount of interface cannot be fabricated at this speed, precision, or resolution by any of the currently available micro-fabrication or printing platforms. It should be noted that flow rates of up to 3 mL min^−1^ could be conveniently achieved using our printing method.

In tissue engineering scenarios, multi-material and multilayer structures are required to mimic the architecture and functionality of real tissues.^[17]^ We also conducted chaotic bioprinting experiments in which we fabricated bands of C2C12 murine skeletal myoblasts within alginate fibers added with GelMA‡. We present the cross-sectional (Figure 5A) and longitudinal view (Figure 5B) of a cell-laden alginate fiber lightly enriched with protein (GelMA) to favor cell proliferation. Well-defined bands of C2C12 cells are distinguished along the hydrogel fibers. Mammalian cells are shear-sensitive; however, the low shear laminar conditions prevalent at the printhead tip enabled high initial cell viabilities (higher than 95%; Figure 5B). Cells survived and proliferated within these fibers. Figure 5C shows cells in a chaotically printed fiber after 7 days of culture. The fibers have been stained to reveal the position of the cell nuclei and their cytoskeletons. Most cells remained within the striations corresponding to the cell-laden ink. After 7 days of culture, the cells began to spread and interact within each other, and some clusters of cells appear (Figure 5C,D). Note that in some cell clusters, multinucleated cells, a signature of myotubule development, started to be evident. After 2 weeks of culture, the proliferating cells elongated while maintaining their initial striation patterns (Figure 5 E,F). This illustrative experiment suggests the potential of chaotic bioprinting to produce massive amounts of living fibers that closely resemble the multilayered structure observed in mammalian tissues.

**Figure 5.**
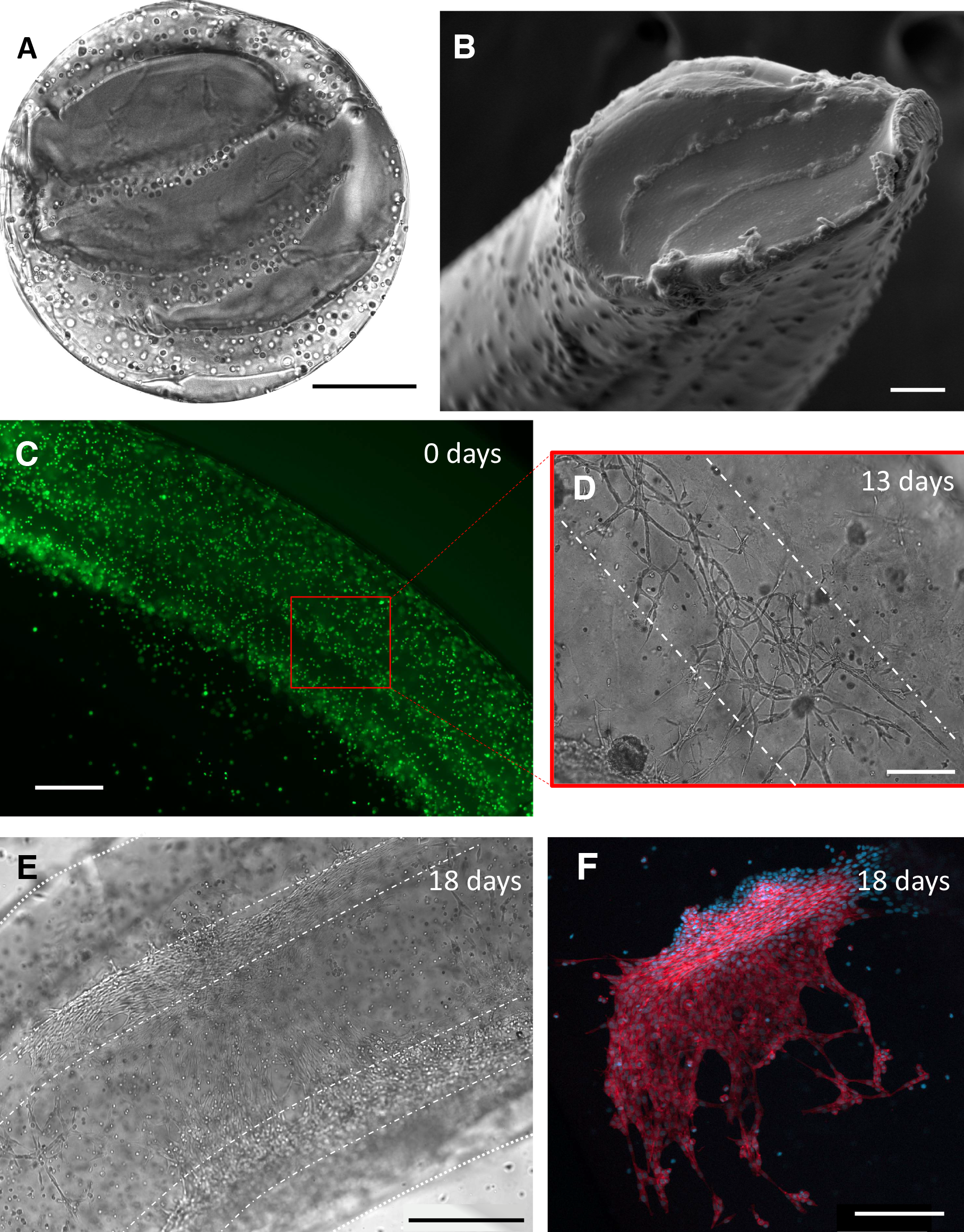
Bioprinting of living micro-tissues: (A) Optical and (B) SEM micrographs of the crosssection view of a construct in which C2C12 cells are chaotically bioprinted in an alginate/GelMA hydrogel using a 3-KSM printhead; Scale bars: 500 μm and 50 μm, respectively. (C) Longitudinal view of a chaotically bioprinted construct; a high cell viability is observed at the initial time, as revealed by a live/dead staining. Scale bar: 500 μm. (D) Cells spread along the chaotically printed striations, preserving their original positions after 13 days of culture. Scale bar: 200 μm. (E) Optical microscopy view of a segment of fiber containing C2C12 cells at 18 days after printing. Scale bar: 500 μm. (F) Close-up of a region stained to reveal F-actin/nuclei, showing the cell spreading and the formation of interacting cell clusters. Cell nuclei can be identified as blue dots. Actin filaments appear in red. Scale bar: 200 μm.

## Conclusion

In this study, we have presented continuous chaotic printing as a strategy that enables full control of the spatial microstructures within a single 3D printed fiber. The key element of this technological platform is the use of an on-line static mixer in the printhead to provide a partial mixing of different materials as they are coextruded through the nozzle tip. In particular, we have adopted the KSM as the first model and have used it to fabricate, in a simple fashion, highly convoluted 3D structures within polymer composites in a continuous stream at high speeds (>1.0 meters of fiber/min). The diameter of the printing head and the number of mixing elements determine the number and thickness of internal lamellae produced according to a process of successive bifurcations that yields an exponential generation of inter-material area. Illustratively, by using 7 internal elements, 128 lamellae of average widths of 7 μm can be generated in a 1 mm cross-section fiber, and an inter-material area of ~67 m^2^ L^−1^ can be achieved. These values for microstructure resolution, internal surface area density, and fabrication speed all exceed the capabilities of any of the currently available commercial microfabrication techniques (i.e., commercial 3D printers) for the creation of microstructure.

The fundamentals underlying chaotic printing are solid, as this type of printing relies on the use of chaotic flows to develop microstructure at an exponential rate in a deterministic manner. Indeed, we have shown that the microstructure resulting from the use of different numbers of KSM elements is amenable to rigorous modeling using CFD simulations, and the resemblance between our experimental and simulation results is remarkable. This precise predictability of the microstructures within a printed construct will greatly expand the application of 3D printing and the complexity of printed composites. A wide spectrum of microstructures can be designed and obtained using this technique. The adoption of different types of static mixer elements (i.e., SMX, Sultzer, and novel *ad hoc* designs), the use of more than two inks (or materials), the manipulation of the injection location, and the dynamic changes in the speed of each one of the injections, can open up possibilities for obtaining structures with various degrees of complexity in a wide range of scales. For instance, we showed that chaotic printing is a simple and versatile micro-and nanofabrication platform and that, when coupled with other fabrication resources, can generate macrostructures with an enormous amount of interfaces between their constituent materials. In an exciting further development, we demonstrated that the output of a continuous 3D chaotic printhead can be fed into an electrospinning nozzle to create fiber meshes with lamellar nanostructures. By doing so, we have shown that the microstructure created by 3D chaotic printing can be further scaled down by three orders of magnitude.

We envision numerous applications of continuous chaotic printing in biomedicine (i.e., bacterial and mammalian cell bioprinting), electronics (i.e., fabrication of high-sensitivity multi-branch electrodes and supercapacitors), and materials science.

## Experimental Section

### Experimental set-up

Our continuous chaotic printer consisted of a syringe pump loaded with two 10-mL disposable syringes, a cylindrical printhead containing from 2 to 7 KSM elements, and a flask containing 550 mL of 1% calcium chloride (Fermont, Productos Químicos Monterrey, Monterrey, NL, Mexico) (Figure 1A). Syringes were loaded with different inks (i.e., particle suspensions in pristine 1% alginate) and connected to one of the two inlet ports located in the lid of the printhead. Details of the geometry of the printer head and the internal KSM elements are shown in Figure 1B,C. The fabrication of printer heads is described in a following subsection (KSM printheads). The syringe pump was set to operate at a flow rate of 0.8 to 1.5 mL min^−1^. We conducted experiments using printheads with different internal diameters, in the range from 5.8 to 2 mm. The tube containing the KSM could be connected to a tip to further reduce the diameter of the final fiber. Tip reducers with an outlet diameter of 4, 2, and 1 mm were used in the experiments presented here (Figure 1D,F,G). The outlet of the tip was submerged in 1% calcium chloride to crosslink the extruded fibers as soon as they exited the tube (Figure 1D).

### KSM printheads

We fabricated our KSM printheads in house. KSM elements were designed using SolidWorks based on the optimum proportions reported in literature.^[49]^ The sets of KSM elements were printed on a P3 Mini Multi Lens 3D printer (EnvisionTEC, Detroit, Michigan) from the ABS Flex White material. We used a length-to-radius ratio of L:3R (Figure 1B). For example, for printheads with an internal diameter of 5.8 mm, the length and diameter of each separate KSM element were 8.7 mm and 5.8 mm, respectively. Sets of 2, 3, 4, 5, 6, and 7 KSM elements, attached to a tube cap, were fabricated to ensure a correct orientation of the ink inlet ports on the cap with respect to the first KSM (Figure 1C). The cap was designed so that each ink inlet was positioned on a different side of the first KSM element to maintain similar initial conditions in all experiments (Figure 1A, C).

### Printing experiments and ink formulations

We used several different ink formulations for the experiments presented here. Inks consisted of particles suspended in 1% alginate or pristine alginate (CAS 9005-38-3, Sigma Aldrich, St. Louis, MO, USA) solutions.

In a first set of experiments, we fabricated fibers loaded with either red or green fluorescent particles. Red and green fluorescent inks were prepared by suspending 1 part of commercial fluorescent particles (Fluor Green 5404 or Fluor Hot Pink 5407; Createx Colors; East Granby, CT, USA) in 9 parts of a 1% aqueous solution of sodium alginate (Sigma Aldrich, St. Louis, MO, USA). The fluorescent particles were previously subjected to three cycles of washing, centrifugation, and decantation to remove surfactants present in the commercial preparation.

We also used chaotic printing to fabricate fibers containing an overall concentration of 0.5% graphite by co-extruding a suspension of 1.0% graphite in alginate solution (1%) and pristine alginate solution (1%) through printheads containing 2, 4, or 6 KSM elements. In addition, we produced control fibers by extruding pristine alginate (without graphite microparticles) through an empty tube, or by co-extruding two streams of ink containing 0.5% graphite microparticles well-mixed by hand in alginate.

In a third set of experiments, we used fluorescent inks based on suspensions of fluorescent *E. coli* bacteria. These fluorescent bacteria were engineered to produce either GFP or RFP. Bacterial inks were prepared by mixing either GFP-or RFP-expressing *E. coli* in 2% alginate solution supplemented with 2% Luria-Bertani (LB) broth (Sigma Aldrich, St. Louis, MO, USA). For ink preparation, bacterial strains were cultivated for 48 h at 37 °C in LB media. Bacterial pellets, recovered by centrifugation, were washed and re-suspended twice in alginate-LB medium. The optical density of the resuspended pellets was adjusted to 0.1 absorbance units before printing (approximately 5 × 10^8^ colony forming units per mL (CFU/mL)). Fibers were printed at a flow rate of 1.5 mL/min and cultured by immersion in LB media for 72 hours. The number of viable cells present in the fibers at different times was determined by conventional plate-counting methods. Briefly, fiber samples of 0.1 g were cultured in tubes containing LB media. The number of viable cells was determined by washing the 0.1 g samples in 1X phosphate-buffered saline (PBS) at pH 7.4 (Gibco, Life Technologies, Carlsbad, CA) to remove the bacteria accumulated in the LB media. Each sample was disaggregated and homogenized in 0.9 mL of PBS. The resultant bacterial suspensions were decimally diluted, seeded onto 1.5% LB-Agar (Sigma Aldrich, St. Louis, MO, USA) plates, and incubated at 37 °C for 36 h.

We also bioprinted muscular murine cells (C2C12 cell line, ATCC CRL 1772) in 1% alginate inks supplemented with 3% gelatin methacryloyl (GelMA) added with a photoinitiator (0.01% LAP). To this purpose, a first ink contained only alginate and GelMA, while the second was cell-laden with C2C12 cells at a concentration of 3×10^6^ cell mL^−1^. Cell laden fibers, obtained by immersion in alginate and then further crosslinked by exposure to UV light at λ=400 nm for 30 s. The bioprinted and cell-laden fibers were immersed in DEMEM culture medium (Gibco, USA) and incubated for 20 days at 37 °C in an 5% CO_2_ atmosphere. Culture medium was renewed every 4th day during the culture period.

In a fifth set of experiments, we produced electrospun nanofiber mats by combining 3D chaotic printing in-line with electrospinning. Fibers produced by 3D chaotic printing were continuously solidified as they were generated by direct feeding into an electrospinning apparatus. In these experiments, we explored two distinct ink pairs for experiments combining 3D continuous chaotic printing and electrospinning. First, we chaotically printed fibers by coextrusion of a pristine alginate ink (4% sodium alginate in water) and polyethylene oxide (7% PEO in water). The resulting PEO-alginate fibers were electrospun (in-line) to produce nanofiber mats.

### Microscopy characterizations

The microstructure of the fibers produced by chaotic printing was analyzed by optical microscopy using an Axio Imager M2 microscope (Zeiss, Germany) equipped with Colibri.2 led illumination and an Apotome.2 system (Zeiss, Germany). Bright-field fluorescence micrographs were used to document the lamellar structures within the longitudinal segments and cross-sections of the fibers. Wide-field images (up to 20 cm^2^) were created using a stitching algorithm included as part of the microscope software (Axio Imager Software, Zeiss, Germany). Fibers were frozen by sudden immersion in liquid nitrogen to facilitate sectioning while preserving the microstructure. The microstructure of the nanofibers produced by chaotic printing coupled with electrospinning was analyzed by atomic force microscopy (AFM) and photo-induced force microscopy (PIFM), a nano-IR technique (Figure S3).

### Mechanical testing of graphite-alginate fibers

We used a universal test bench machine (Tinus Olsen h10kn; PA; USA), with a load cell of 50 N at a rate of 35 mm min^−1^, to evaluate the mechanical properties of alginate fibers containing 0.5% graphite particles and produced by different printing strategies. Specifically, we conducted tensile testing on fibers produced from a mixture of 0.5% graphite microparticles in alginate, generated either by random mixing and extrusion through an empty pipe (a control without lamellar structures) or by continuous chaotic printing using 2, 4, or 6 KSM elements. In these experiments, the gauge length between clamps was set to 25 mm. Stress-strain curves were obtained for each of the five different formulations. We determined the maximum tensile strength, strain at break, and Young modulus of the fibers from stress-strain data.

### Computational simulations

The system was simulated using a finite element model (FEM) strategy in COMSOL Multiphysics 5. First, a 3D model was designed and solved, using laminar flow equations and a stationary solver, to determine the velocity field in the system for the various experimental scenarios explored. A fluid viscosity value of 1P and a density of 1000 kg/m^3^ were used. A time dependent solver was then used to track up to 10^5^ massless particles using particle tracking for fluid flow physics in the previously solved stationary velocity field. The simulation was discretized with a reasonable fine mesh composed of free triangular elements. Mesh sensitivity studies were conducted to ensure the consistency of results. No-slip boundary conditions were imposed in the fluid flow simulation, while a freeze boundary condition was employed for the particle tracing module. The interface length was determined by importing the output results from the cross-section of the fibers (a set of points describing the interface position) into CorelDraw software X5 (Corel Corporation, Canada), drawing Bezier curves over the striations, and establishing the length of the curves using the software (Figure S6).

## Supporting information

Electronic Supplementary Information

## Acknowledgements

CCM and MDdLD contributed equally to this work. GTdS acknowledges the funding received from CONACyT (Consejo Nacional de Ciencia y Tecnología, México) (66839), L’Oréal-UNESCO-CONACyT-AMC (National Fellowship for Women in Science, Mexico) and UC-MEXUS. MMA and GTdS acknowledge funding provided from CONACyT, Fronteras de la Ciencia No. 2442. Yu Shrike Zhang acknowledges the funding by the National Institutes of Health (R21EB025270, R21EB026175, R00CA201603, R01EB028143) and the Brigham Research Institute. AK would like to acknowledge funding from the National Institutes of Health (HL137193, 5R01AR057837, 1R01EB021857). This research has been partially funded by the Tecnológico de Monterrey and the Massachusetts Institute of Technology (MIT) Nanotechnology Program. HKW and SOMC greatly acknowledge funding provided by Cátedra Federico Baur, Tecnológico de Monterrey. We gratefully acknowledge the experimental assistance of Gyan Prakash, Everardo González-González, Aimé Alexandra Cuellar-Monterrubio, Alan Roberto Márquez-Ipiña, Sara Cristina Pedroza, Felipe López-Pacheco, Matías Lobo-Zegers, Zamantha Escobedo-Avellaneda, and Esther Pérez-Carrillo. We acknowledge the valuable assistance of Centro de Investigación en Química Aplicada (CIQA) in Saltillo, Coahuila, México.

## Author Contributions

GTdS, MMA, designed the study. CChM and MDdLD developed the experimental set-up and conducted most of the printing experiments. MS performed all the computational simulations. CCMB and JFYdL fabricated the printheads by stereolitographic 3D printing. CFCG, JEBM and CChM. conducted most of the bioprinting experiments. SH, NAGF, and CChM performed the electrospinning experiments. MAA conducted PiMF characterization experiments. IGG and CChM conducted the mechanical testing experiments. GTdS and MMA produced the first complete draft of the manuscript. GTdS, MMA, MM, YSZ, CChM, SOMCh, CAR, HKW, and AK significantly edited the manuscript. CChM, GTdS, MMA, MDdLD, MS, and MAA prepared illustrations. All authors read, commented, and approved the manuscript.

‡ Methacryloyl-gelatin (GelMA) is a UV cross-linkable material that has been widely used in tissue engineering applications; it contains cell binding domains, it is biodegradable, and amenable to microfabrication.

## References

[1] L. Sheng, J. Chang, L. Jiang, Z. Jiang, Z. Liu, T. Wei, Z. Fan, Adv. Funct. Mater. 2018, 28, 1800597.

[2] X. Shen, Z. Wang, Y. Wu, X. Liu, Y. B. He, Q. Zheng, Q. H. Yang, F. Kang, J. K. Kim, Mater. Horizons 2018, 5, 275.

[3] H. An, T. Habib, S. Shah, H. Gao, M. Radovic, M. J. Green, J. L. Lutkenhaus, Sci. Adv. 2018, 4, eaaq0118.

[4] H. L. Gao, Y. B. Zhu, L. B. Mao, F. C. Wang, X. S. Luo, Y. Y. Liu, Y. Lu, Z. Pan, J. Ge, W. Shen, Y. R. Zheng, L. Xu, L. J. Wang, W. H. Xu, H. A. Wu, S. H. Yu, Nat. Commun. 2016, 7, 12920.

[5] P. He, J. R. Brent, H. Ding, J. Yang, D. J. Lewis, P. O’Brien, B. Derby, Nanoscale 2018, 10, 5599.

[6] I. J. Chung, W. Kim, W. Jang, H. W. Park, A. Sohn, K. B. Chung, D. W. Kim, D. Choi, Y. T. Park, J. Mater. Chem. A 2018, 6, 3108.

[7] A. Goyanes, J. Wang, A. Buanz, R. Martínez-Pacheco, R. Telford, S. Gaisford, A. W. Basit, Mol. Pharm. 2015, 12, 4077.

[8] D. B. Kolesky, R. L. Truby, A. S. Gladman, T. A. Busbee, K. A. Homan, J. A. Lewis, Adv. Mater. 2014, 26, 3124.

[9] H. W. Kang, S. J. Lee, I. K. Ko, C. Kengla, J. J. Yoo, A. Atala, Nat. Biotechnol. 2016, 34, 312.

[10] A. K. Miri, I. Mirzaee, S. Hassan, S. Mesbah Oskui, D. Nieto, A. Khademhosseini, Y. S. Zhang, Lab Chip 2019.

[11] A. K. Miri, A. Khalilpour, B. Cecen, S. Maharjan, S. R. Shin, A. Khademhosseini, Biomaterials 2019, 198, 204.

[12] A. L. Rutz, K. E. Hyland, A. E. Jakus, W. R. Burghardt, R. N. Shah, Adv. Mater. 2015, 27, 1607.

[13] Q. Ge, A. H. Sakhaei, H. Lee, C. K. Dunn, N. X. Fang, M. L. Dunn, Sci. Rep. 2016, 6, 31110.

[14] J. O. Hardin, T. J. Ober, A. D. Valentine, J. A. Lewis, Adv. Mater. 2015, 27, 3279.

[15] W. Liu, Y. S. Zhang, M. A. Heinrich, F. De Ferrari, H. L. Jang, S. M. Bakht, M. M. Alvarez, J. Yang, Y.-C. Li, G. Trujillo-de Santiago, A. K. Miri, K. Zhu, P. Khoshakhlagh, G. Prakash, H. Cheng, X. Guan, Z. Zhong, J. Ju, G. H. Zhu, X. Jin, S. R. Shin, M. R. Dokmeci, A. Khademhosseini, Adv. Mater. 2017, 29, 1604630.

[16] D. Kang, G. Ahn, D. Kim, H. W. Kang, S. Yun, W. S. Yun, J. H. Shim, S. Jin, Biofabrication 2018, 10, 035008.

[17] J. U. Lind, T. A. Busbee, A. D. Valentine, F. S. Pasqualini, H. Yuan, M. Yadid, S. J. Park, A. Kotikian, A. P. Nesmith, P. H. Campbell, J. J. Vlassak, J. A. Lewis, K. K. Parker, Nat. Mater. 2017, 16, 303.

[18] Q. Pi, S. Maharjan, X. Yan, X. Liu, B. Singh, A. M. van Genderen, F. Robledo-Padilla, R. Parra-Saldivar, N. Hu, W. Jia, C. Xu, J. Kang, S. Hassan, H. Cheng, X. Hou, A. Khademhosseini, Y. S. Zhang, Adv. Mater. 2018, 1706913.

[19] M. M. Alvarez-Hernández, T. Shinbrot, J. Zalc, F. J. Muzzio, Chem. Eng. Sci. 2002, 57, 3749.

[20] P. E. Arratia, T. Shinbrot, M. M. Alvarez, F. J. Muzzio, Phys. Rev. Lett. 2005, 94, 084501.

[21] T. J. Ober, D. Foresti, J. A. Lewis, Proc. Natl. Acad. Sci. U. S. A. 2015, 112, 12293.

[22] B. Grigoryan, S. J. Paulsen, D. C. Corbett, D. W. Sazer, C. L. Fortin, A. J. Zaita, P. T. Greenfield, N. J. Calafat, J. P. Gounley, A. H. Ta, F. Johansson, A. Randles, J. E. Rosenkrantz, J. D. Louis-Rosenberg, P. A. Galie, K. R. Stevens, J. S. Miller, Science (80-.). 2019, 364, 458.

[23] D. M. Hobbs, M. M. Alvarez, F. J. Muzzio, Fractals 1997, 05, 395.

[24] F. J. Muzzio, M. M. Alvarez, S. Cerbelli, Massimiliano Giona, A. Adrover, Chem. Eng. Sci. 2000, 55, 1497.

[25] M. M. Alvarez, F. J. Muzzio, S. Cerbelli, A. Adrover, M. Giona, Phys. Rev. Lett. 1998, 81, 3395.

[26] M. M. Alvarez, J. M. Zalc, T. Shinbrot, P. E. Arratia, F. J. Muzzio, AIChE J. 2002, 48, 2135.

[27] G. Trujillo-de Santiago, M. M. Alvarez, M. Samandari, G. Prakash, G. Chandrabhatla, P. I. Rellstab-Sánchez, B. Byambaa, P. Pour Shahid Saeed Abadi, S. Mandla, R. K. Avery, A. Vallejo-Arroyo, A. Nasajpour, N. Annabi, Y. S. Zhang, A. Khademhosseini, Mater. Horizons 2018, 5, 813.

[28] D. M. Hobbs, F. J. Muzzio, Chem. Eng. J. 1997, 67, 153.

[29] J. M. Zalc, E. S. Szalai, F. J. Muzzio, S. Jaffer, AIChE J. 2002, 48, 427.

[30] D. S. Kim, S. H. Lee, T. H. Kwon, C. H. Ahn, Lab Chip 2005, 5, 739.

[31] C. Y. Lee, W. T. Wang, C. C. Liu, L. M. Fu, Chem. Eng. J. 2016, 288, 146.

[32] D. Nowak, W. Morrison, H. K. Wickramasinghe, J. Jahng, E. Potma, L. Wan, R. Ruiz, T. R. Albrecht, K. Schmidt, J. Frommer, D. P. Sanders, S. Park, Sci. Adv. 2016, 2, e1501571.

[33] D. Choudhury, S. Anand, M. W. Naing, Int. J. Bioprinting 2018, 4, 139.

[34] A. N. Leberfinger, S. Dinda, Y. Wu, S. V. Koduru, V. Ozbolat, D. J. Ravnic, I. T. Ozbolat, Acta Biomater. 2019, 95, 32.

[35] M. A. Heinrich, W. Liu, A. Jimenez, J. Yang, A. Akpek, X. Liu, Q. Pi, X. Mu, N. Hu, R. M. Schiffelers, J. Prakash, J. Xie, Y. S. Zhang, Small 2019, 15, 1805510.

[36] A. Adrover, M. Giona, F. J. Muzzio, S. Cerbelli, M. M. Alvarez, Phys. Rev. E 1998, 58, 447.

[37] D. M. Hobbs, F. J. Muzzio, Chem. Eng. Sci. 1998, 53, 3199.

[38] D. M. Hobbs, P. D. Swanson, F. J. Muzzio, Chem. Eng. Sci. 1998, 53, 1565.

[39] A. S. Gladman, E. A. Matsumoto, R. G. Nuzzo, L. Mahadevan, J. A. Lewis, Nat. Mater. 2016, 15, 413.

[40] M. Schaffner, P. A. Rühs, F. Coulter, S. Kilcher, A. R. Studart, Sci. Adv. 2017, 3, eaao6804.

[41] R. M. Stubbendieck, C. Vargas-Bautista, P. D. Straight, Front. Microbiol. 2016, 7, 1234.

[42] G. Bodelón, V. Montes-García, C. Costas, I. Pérez-Juste, J. Pérez-Juste, I. Pastoriza-Santos, L. M. Liz-Marzán, ACS Nano 2017, 11, 4631.

[43] R. M. Stubbendieck, P. D. Straight, J. Bacteriol. 2016, 198, 2145.

[44] M. Whiteley, S. P. Diggle, E. P. Greenberg, Nature 2017, 551, 313.

[45] S. Alberghini, E. Polone, V. Corich, M. Carlot, F. Seno, A. Trovato, A. Squartini, FEMS Microbiol. Lett. 2009, 292, 149.

[46] J. L. Connell, J. Kim, J. B. Shear, A. J. Bard, M. Whiteley, Proc. Natl. Acad. Sci. 2014, 111, 18255.

[47] S. E. Darch, O. Simoska, M. Fitzpatrick, J. P. Barraza, K. J. Stevenson, R. T. Bonnecaze, J. B. Shear, M. Whiteley, Proc. Natl. Acad. Sci. U. S. A. 2018, 115, 4779.

[48] A. Bohle, B. Aeikens, A. Eenboom, L. Fronholt, W. R. Plate, J.-C. Xiao, A. Greschniok, M. Wehrmann, Kidney Int. 1998, 54, S186.

[49] O. Byrde, M. L. Sawley, Chem. Eng. J. 1999, 72, 163.

