## Supplementary material for "Using Chaos for Facile High-throughput Fabrication of Ordered Multilayer Micro- and Nanostructures": Electronic Supplementary Information

*Carolina Chavez-Madero^#^, Maria Diaz de Leon-Derby^#^, Mohamadmahdi Samandari, Christian Carlos Mendoza-Buenrostro, Carlos Fernando Ceballos-Gonzalez, Edna Johana Bolivar-Monsalve, Sunshine Holmberg, Norma Alicia Garza-Flores, Mohammad Ali Almajhadi, Ivonne Gonzalez-Gamboa, Juan Felipe Yee-de Leon, Sergio Omar Martinez-Chapa, Ciro Angel Rodriguez-Gonzalez, Kumar Wickramasinghe, Marc Madou, Ali Khademhosseini, Yu Shrike Zhang, Mario Moises Alvarez^*^, Grissel Trujillo-de Santiago*^*^

#These authors contributed equally to this work

**Supplementary Text and figures**

In Figure S1, we present results from a series of printing experiments where two alginate inks containing suspended fluorescent particles (red and green, respectively) were coextruded through a nozzle that contained 2 to 6 KSM elements. Figure S1 shows (a) the cross-sectional and (a,b) longitudinal views of the printed alginate fibers. Experimental and Computational results are presented. Filaments with different degrees of intimacy in their structures, as observed by fluorescence microscopy, could be fabricated. Simulation results, obtained by solving the Navier-Stoke equations of fluid motion using computational fluid dynamics (CFD)^[37,38]^, closely predicted the features of the microstructure resulting from a given set of experimental conditions (i.e., the number of inks, number of elements, and flow rate ratios). In Figure S1c, we estimate the number of striations (s) and the average distance between striations (δ), resulting from the use of printing heads containing an increasing number of KSM-elements.

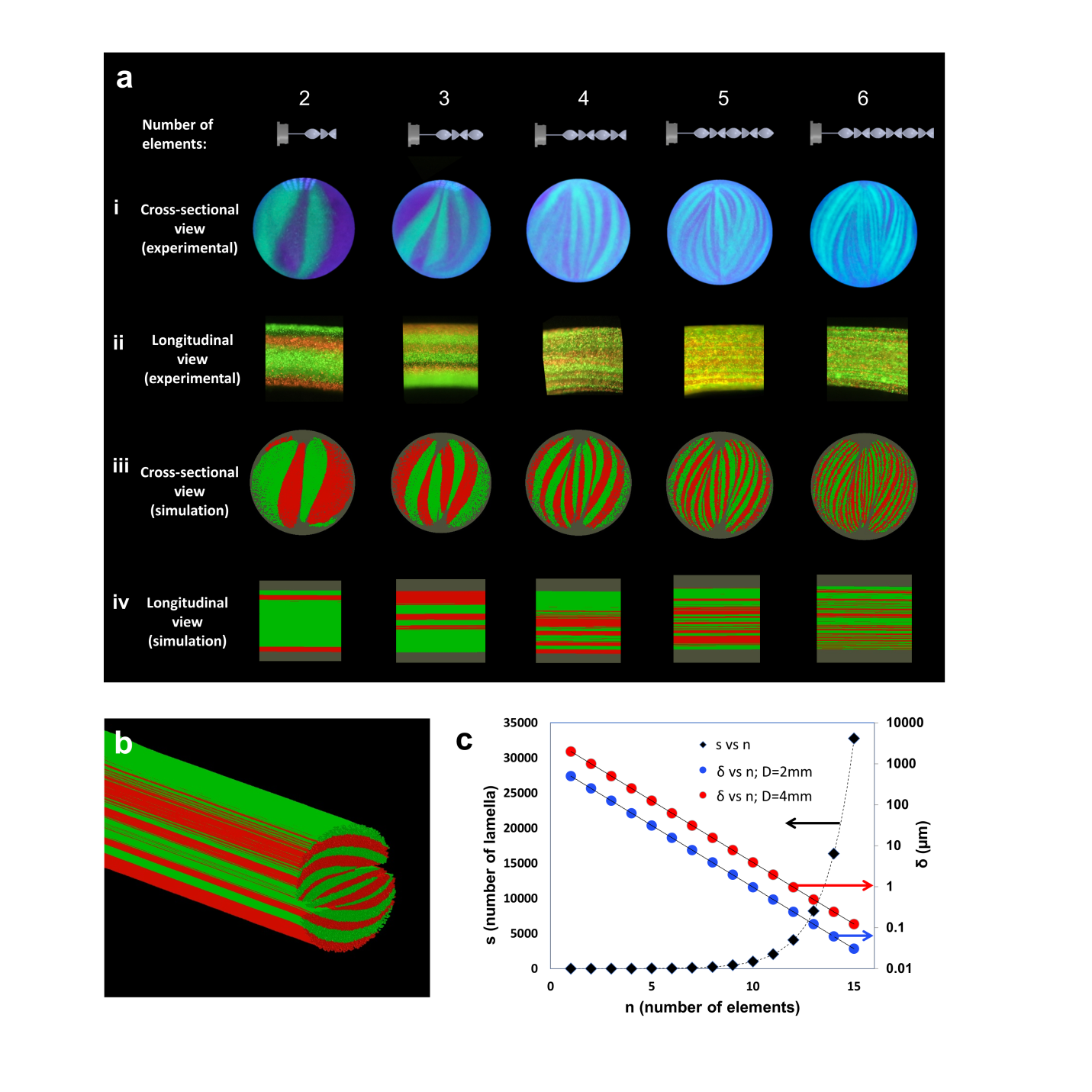

**Figure S1.** Creation of high-resolution microstructure using chaotic printing. (a) The co-extrusion of two different inks, consisting of particle-laden (red and green) alginate, yielded a complex lamellar structure. (i) Cross-section and (ii) lengthwise views of alginate fibers printed with a 2-, 3-, 4-, 5-, and 6‑element KSM using two fluorescent inks (i, ii). Computational simulation of the (iii) cross-section and (iv) length-wise microstructures obtained after mixing two types of massless particles (green and red) using 2, 3, 4, 5, and 6 KSM elements (iii, iv). (b) 3-D view of computational recreation of the microstructures obtained in a segment of fiber (longitudinal and cross-sectional views) after mixing two types of massless particles using 4 KSM elements. (c) Number of striations (s) achieved when using certain number of mixing elements in nozzles with different diameters. The resolution, namely the number of lamellae and the distance between them (δ), can be tuned using different numbers of KSM elements. For small-diameter extruders, nanometer-length scales could be achieved if more than 10 elements were used.

As a general requirement for printability in our chaotic printing setup, inks should behave as Newtonian fluids at the velocity of injection, and exhibit compatibility in terms of their chemical nature (i.e., density, hydrophobicity/hydrophilicity, interface tension, and viscosity). In principle, a wide portfolio of Newtonian cross-linkable inks can be used for chaotic printing. Figure S2 shows the shear stress and effective viscosity at different strain rates of the alginate inks used in our experiments.

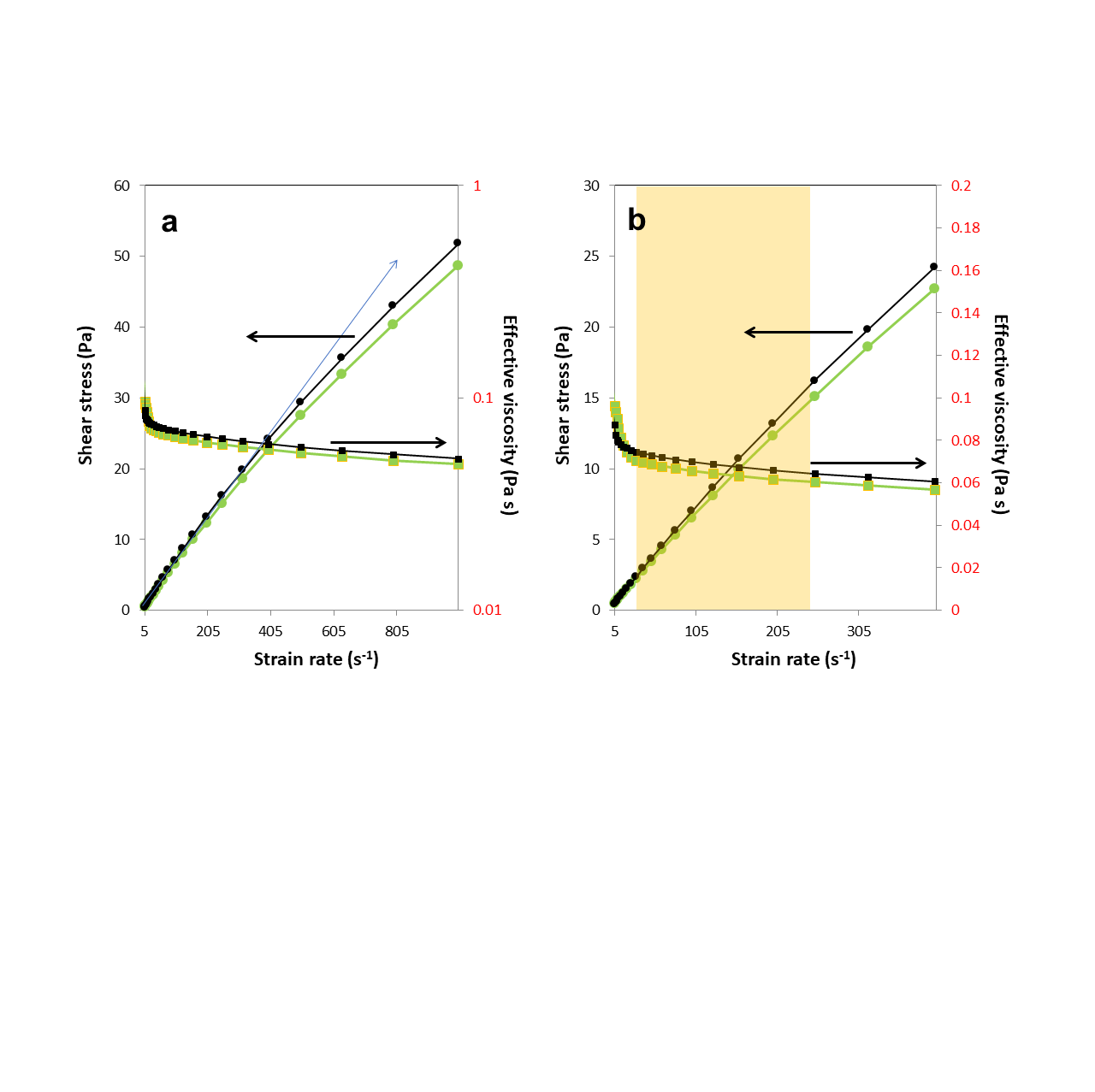

**Figure S2.** (a) Viscosity of an alginate-based ink composed of 0.5% graphite microparticles suspended in a 1% alginate solution (black line) and an alginate-based bio-ink composed of *Escherichia coli* bacteria (~10^8^ bacteria/mL) suspended in a 1% alginate solution (green line). (b) Close-up of the region between strain rate values of 5 to 400 s^-1^. The yellow shaded area indicates the range of strain rates relevant to our chaotic printing experiments.

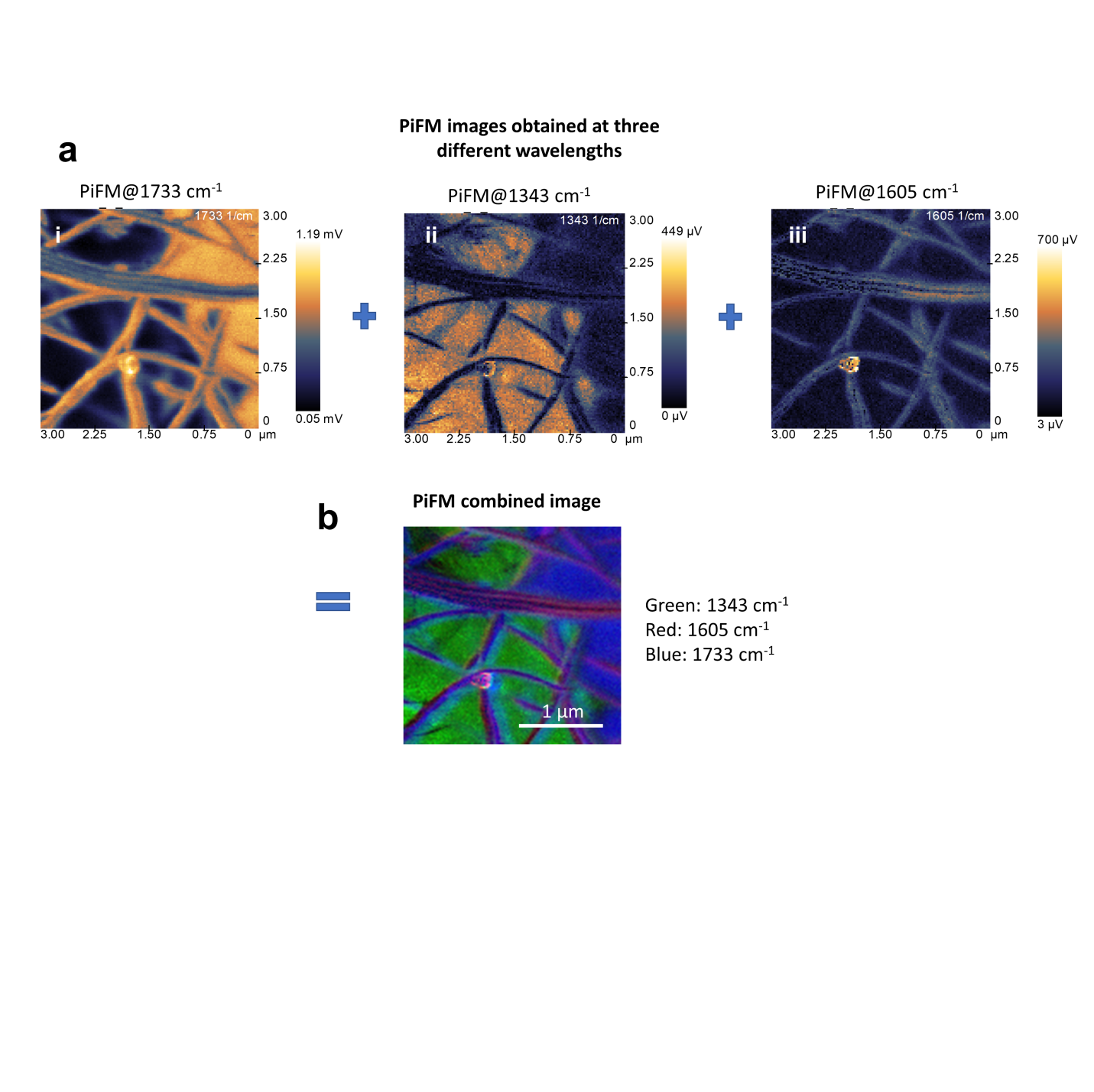

**Figure S3.** Visualization of the lamellar nanostructure in nanospun fibers developed by combining continuous chaotic printing and electrospinning. Micrographs shown in Fig. 5b (i-iv) were obtained by first (a) obtaining photo-induced force microscopy (PiFM) images at different wave numbers (wavelengths): (i) 1733 cm^-1^; (ii) 1343 cm^-1^; (iii) 1605 cm^-1^, and then (b) overlapping all of them into a single micrograph.

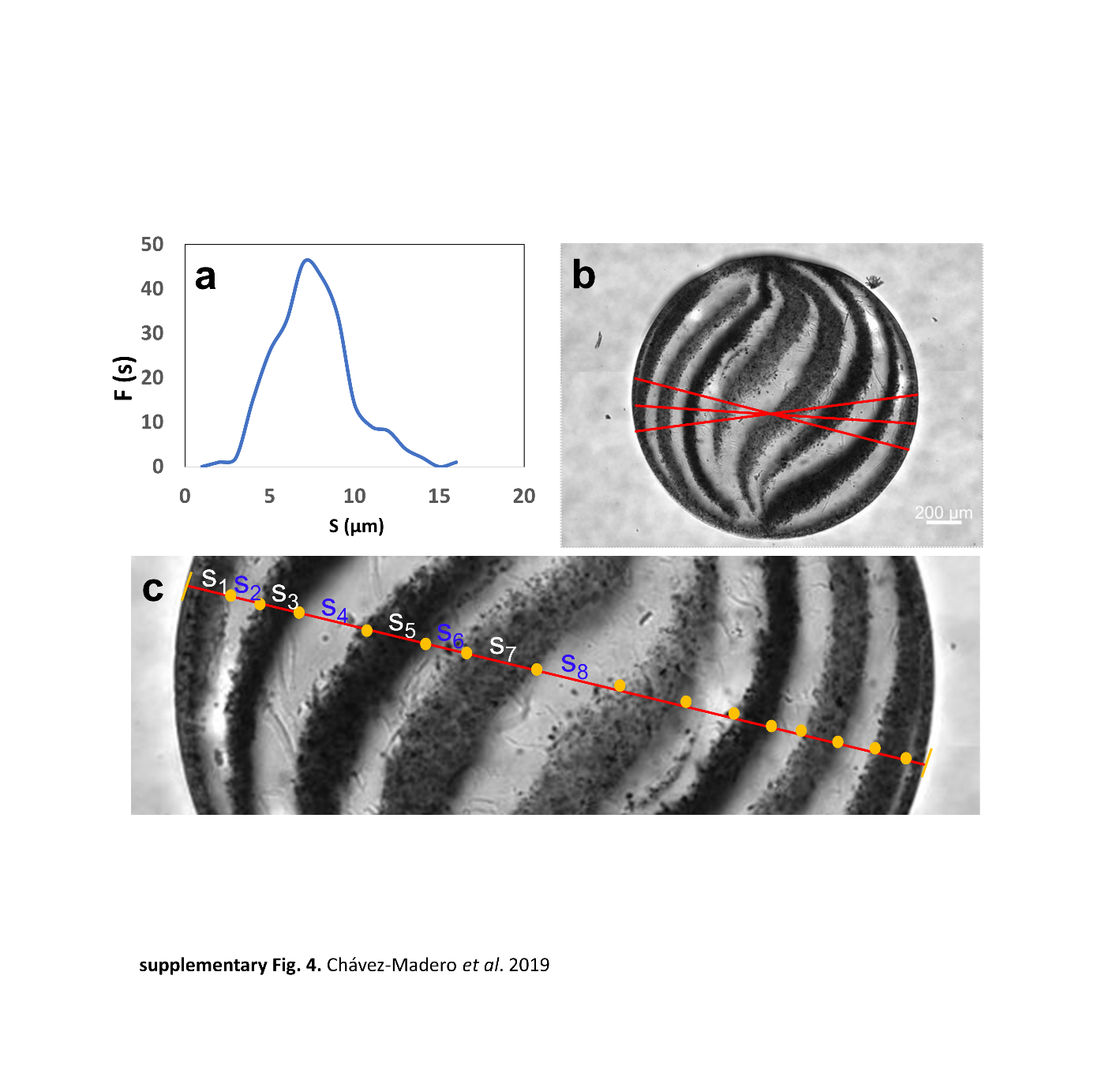

**Figure S4.** Determination of striation thickness distribution (STD) in chaotically printed constructs. To calculate the (a) STD characteristic of a chaotically printed construct, (b) a family of straight centerlines (marked in red) was traced in a high-resolution image of the construct at slightly different angles. (c) Lamella thicknesses were calculated along each centerline (s_i_) using image analysis techniques. The length of the red line is 2 mm.

Figure S5 presents a mechanical characterization of hybrid graphite-alginate fibers fabricated either by extrusion through an empty tube or by continuous chaotic printing. In all cases, the resulting fibers contained 0.5% graphite microparticles in a 1% sodium alginate matrix. Fibers were crosslinked by extrusion into a calcium chloride solution.

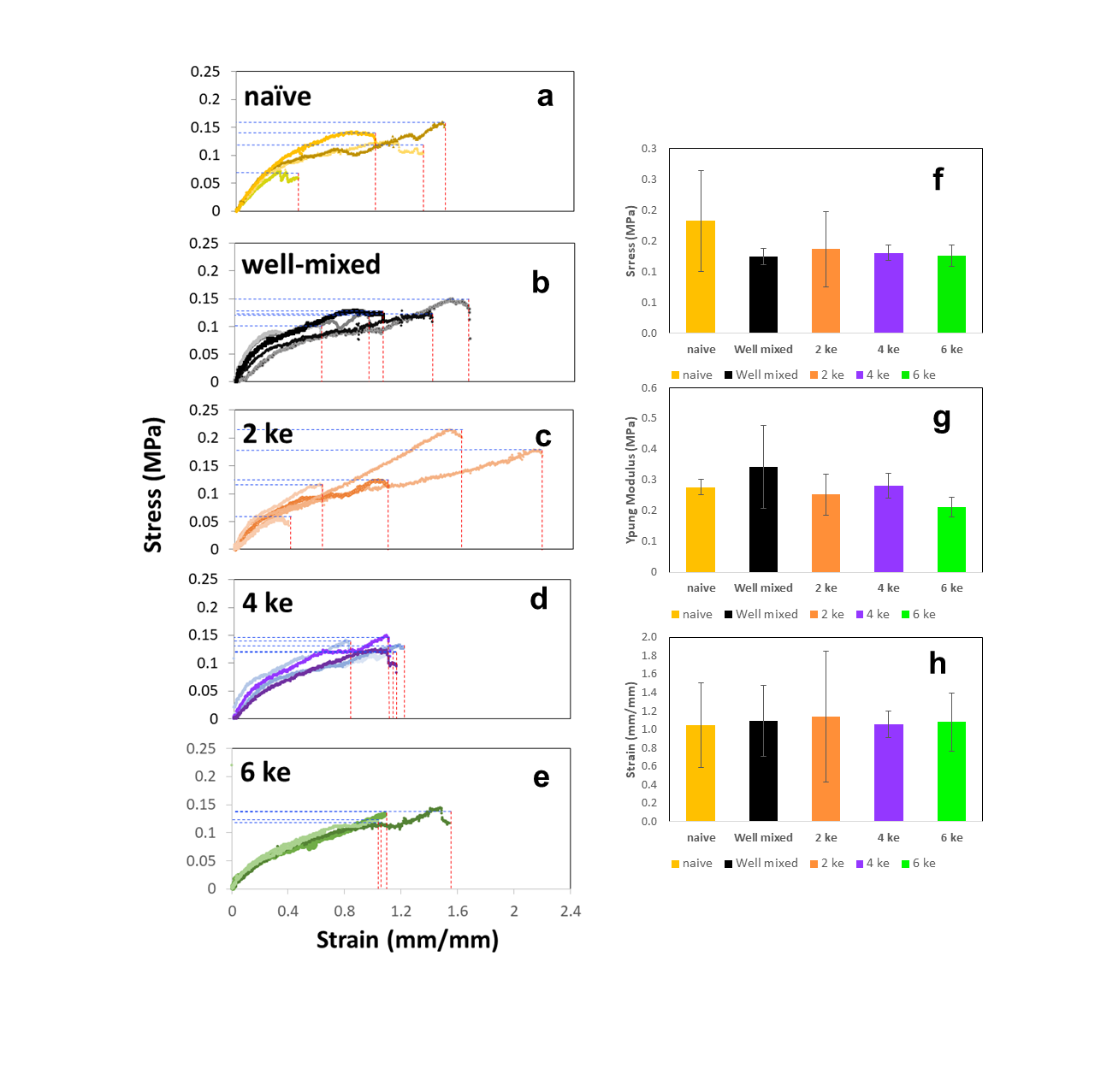

**Figure S5.** Stress versus strain curves for fibers fabricated by extrusion of a well-mixed ink through a tube (internal diameter = 2mm) without internal KSM elements: (a) pristine alginate fiber (1% sodium alginate in water), and (b) a hybrid fiber composed of 0.25% graphite microparticles “well-mixed” in a 1% sodium alginate solution. (C-E) Fibers fabricated by co-extruding a naïve alginate solution (1% sodium alginate in water) and a suspension of graphite-microparticles in alginate (0.5% graphite microparticles in a 1% sodium alginate solution) through a printhead containing (c) 2, (d) 4, (e) and 6 KSM elements. (f) Stress at break, (g) elastic modulus, and (h) maximum strain of fibers fabricated by different strategies: extrusion of naïve alginate (yellow bar) through an empty pipe; extrusion of a well-mixed 0.25% graphite suspension in alginate (black bar); and chaotic printing of naïve alginate and a 0.5% graphite suspension in alginate through a printhead containing two (orange bar), four (purple bar), and six (yellow bar) KSM elements.

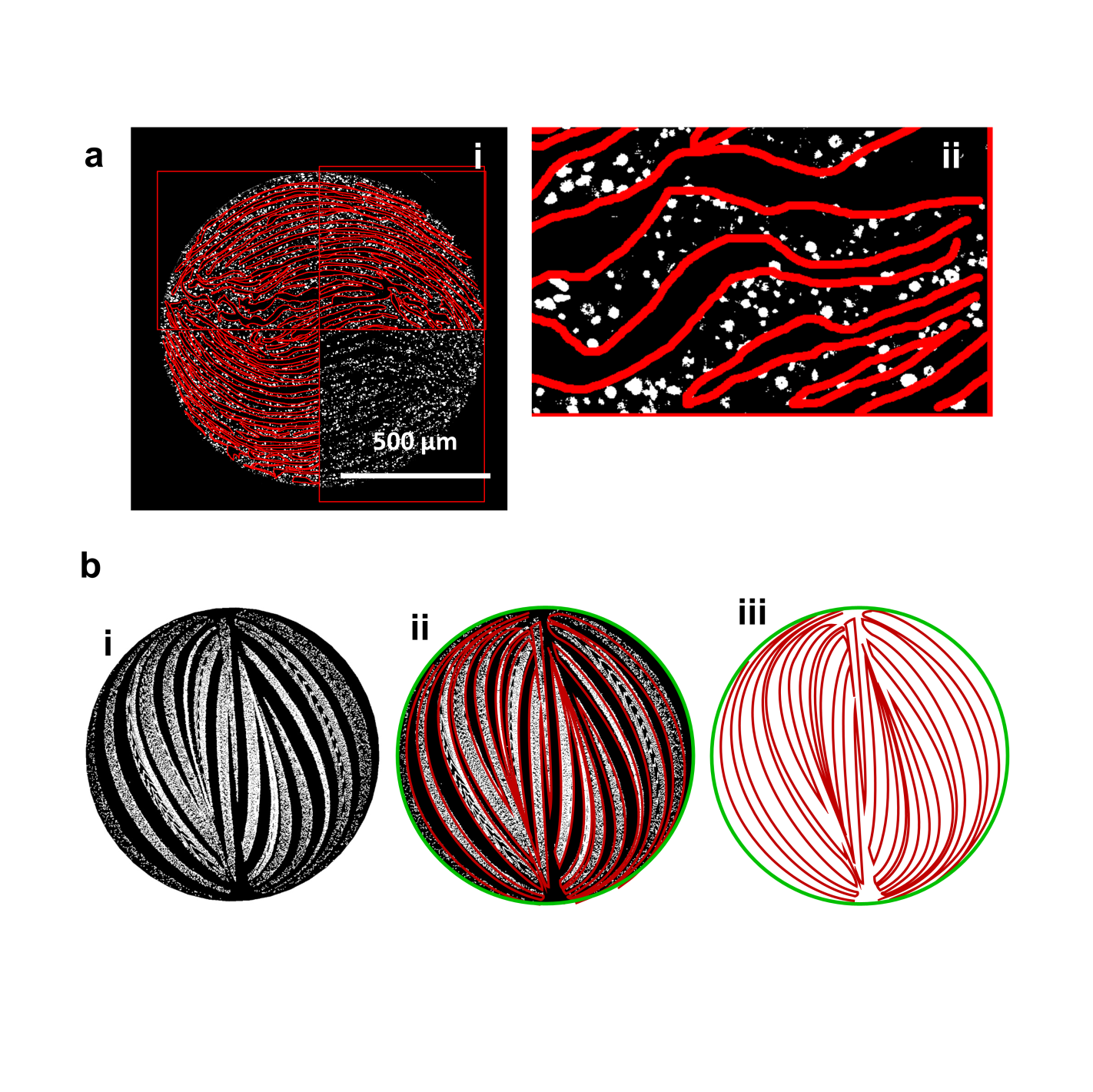

**Figure S6.** Determination of the total interface in chaotically printed constructs. (a) Experimental determination of the interface between zones populated by bacteria and naïve alginate areas. (i) Lines following the interface (bordering each populated lamella) were traced, and (ii) the lengths of those lines were summed using image analysis techniques. The ratio between the total amount of accumulated inter-lamellar interface (*L*) and the perimeter of the fiber (p) was then calculated. (b) Computational fluid dynamics (CFD) determination of the (*L*/p) ratio at the cross-section of chaotically printed using constructs. (i) The interface shared between two co-extruded materials was recreated by tracing the deformation of an interface composed of 100,000 discrete points through repeated splitting and reorientation cycles along the KSM elements in the printheads. (ii) To calculate the total length of the interface (*L*), Bezier curves were manually interpolated over those points along the interface (red), and the perimeter (green) using CorelDraw software, and (iii) the length of the interfaces (red) and the perimeter (green) were determined using the built-in function “object properties” of CorelDraw.
